# The Salt-Inducible Kinase inhibitor YKL-05-099 suppresses MEF2C function and acute myeloid leukemia progression *in vivo*

**DOI:** 10.1101/636969

**Authors:** Yusuke Tarumoto, Shan Lin, Jinhua Wang, Joseph P. Milazzo, Yali Xu, Nathanael S. Gray, Kimberly Stegmaier, Christopher R. Vakoc

## Abstract

Lineage-defining transcription factors (TFs) are compelling targets for leukemia therapy, yet they are among the most challenging proteins to modulate directly with small molecules. We previously used CRISPR screening to identify a Salt-Inducible Kinase 3 (SIK3) requirement for the growth of acute myeloid leukemia (AML) cell lines that overexpress the lineage TF MEF2C. In this context, SIK3 maintains MEF2C function by directly phosphorylating histone deacetylase 4 (HDAC4), a repressive cofactor of MEF2C. Here, we evaluated whether inhibition of SIK3 with the tool compound YKL-05-099 can suppress MEF2C function and attenuate disease progression in animal models of AML. Genetic targeting of SIK3 or MEF2C selectively suppressed the growth of transformed hematopoietic cells under *in vitro* and *in vivo* conditions. Similar phenotypes were obtained when exposing cells to YKL-05-099, which caused cell cycle arrest and apoptosis in MEF2C-expressing AML cell lines. An epigenomic analysis revealed that YKL-05-099 rapidly suppressed MEF2C function by altering the phosphorylation state and nuclear localization of HDAC4. Using a gatekeeper allele of *SIK3*, we found that the anti-proliferative effects of YKL-05-099 occurred through on-target inhibition of SIK3 kinase activity. Based on these findings, we treated two different mouse models of MLL-AF9 AML with YKL-05-099, which attenuated disease progression *in vivo* and extended animal survival at well-tolerated doses. These findings validate SIK3 as a therapeutic target in MEF2C-positive AML and provide a rationale for developing drug-like inhibitors of SIK3 for definitive pre-clinical investigation and for studies in human patients with leukemia.

**Key Points:** AML cells are uniquely sensitive to genetic or chemical inhibition of Salt-Inducible Kinase 3 *in vitro* and *in vivo*.

A SIK inhibitor YKL-05-099 suppresses MEF2C function and AML *in vivo*.

## Introduction

Acute myeloid leukemia (AML) is a hematopoietic malignancy characterized by an aberrant self-renewal potential of myeloid progenitor cells. This cellular phenotype is imposed by a diverse set of genetic drivers, which often promote leukemogenesis via direct or indirect deregulation of lineage-defining transcription factors (TFs)^1–6^. Hence, a reprogrammed transcriptional state of cell identity underpins that pathogenesis of AML, a process that is known to render leukemia cells hypersensitive to perturbations of lineage TFs when compared to normal myeloid progenitors^6^. While lineage TFs represent an important category of molecular vulnerabilities in AML, this finding has yet to achieve clinical significance since TFs are among the most challenging proteins to target directly with drugs^7,8^. For this reason, a major objective in AML research is in identifying chemical matter capable of direct or indirect TF modulation.

Myocyte enhancer factor 2C (MEF2C) is a leukemogenic TF that regulates normal cell fate specification programs, including myogenesis, neurogenesis, and hematopoiesis^9^. Knockout studies in mice indicate that *Mef2c* is essential in the normal lymphoid and megakaryocytic lineages, but is largely dispensable for myelopoiesis and for hematopoietic stem cell self-renewal^10–13^. Insertional mutagenesis screens performed in mice first revealed a leukemogenic function of MEF2C^14^, which was later shown to be overexpressed in a variety of human myeloid and lymphoid cancers in association with poor clinical outcomes^15–21^. The *MLL*-rearranged subtypes of leukemia exemplify a group of leukemias with high MEF2C expression. In this disease, MLL fusion oncoproteins induce an active chromatin state and transcriptional hyper-activation at the *MEF2C* locus^9,15,16^. This results in overexpression of MEF2C, which promotes enhancer-mediated gene activation to promote self-renewal, tissue invasion, and chemotherapy resistance^15,16,20,22^. Importantly, it has been shown that MLL fusion AML cells are addicted to continuous MEF2C expression for their growth and viability^15,22^. The powerful nature of MEF2C addiction in *MLL*-rearranged AML has been most convincingly demonstrated in the hypomorphic *Mef2c^S222A/S222A^* mouse strain, which lacks any detectable developmental abnormalities, but is entirely resistant to leukemic transformation by the MLL-AF9 oncoprotein^21^. Collectively, these genetic experiments validate MEF2C as a vulnerability in AML cells and an attractive target for therapy.

The transcriptional output of MEF2C is dynamically regulated during cell differentiation by several kinase signaling cascades^9^, which presents a strategy for pharmacological MEF2C modulation in cancer. For example, kinases are known to regulate the interaction between MEF2C and the class IIa family of histone deacetylases (HDAC4, HDAC5, HDAC7, and HDAC9)^23,24^, which bind directly to the MADS box/MEF2 domain of MEF2C to form a complex on DNA that is incapable of transcriptional activation^25,26^. Each class IIa HDAC can be phosphorylated by several different kinases, such as calmodulin-dependent protein kinase (CaMK) and Salt-Inducible Kinases (SIKs), at conserved serine residues to promote their interaction with 14-3-3 proteins, which function to sequester HDAC proteins in the cytoplasm^23,27,28^. In addition, MEF2C can be directly phosphorylated by MARK kinases at S222 to promote its transcriptional function^21^. Through such mechanisms, kinase signaling pathways are able to control MEF2C function in a variety of cellular contexts^23,24,27^.

We previously applied kinase domain-focused CRISPR screening to human cancer cell lines in search of context-specific dependencies, which revealed a correlation between Salt-Inducible Kinases (SIK3, in a partially redundant manner with SIK2) and MEF2C essentiality in AML^22^. Our subsequent mechanistic experiments showed that inactivation of SIK3 induced the formation of HDAC4-MEF2C complexes at enhancer DNA elements. This triggered a reduction in vicinal histone lysine acetylation and transcriptional suppression of MEF2C target genes^22^. This study demonstrated a mechanistic link between SIK3 and MEF2C in AML and raised the hypothesis that pharmacological targeting of SIK3 might have therapeutic significance in this disease. Here, we tested this hypothesis using the tool compound YKL-05-099, which inhibits the SIK kinase family and has a suitable bioavailability for pre-clinical studies in mice^29^. As described below, our experiments revealed that YKL-05-099 suppresses the transcriptional output of MEF2C and attenuates disease progression in animal models of *MLL*-rearranged AML.

## Materials and Methods

### Plasmid constructions

The lentiviral sgRNA expression vectors (LRG2.1, Addgene_108098; LRG, Addgene_65656) and the lentiviral Cas9 (LentiV_Cas9_puro, Addgene_108100) or luciferase (Lenti-luciferase-P2A-Neo, Addgene_105621) expression vectors were described previously^22,30^. LentiV_Cas9_Blast (Addgene_125592) was derived from LentiV_Cas9_puro by replacing a puromycin resistance gene with a blasticidin resistance gene. LRG2.1_Neo (Addgene_125593) or LRG2.1_Puro (Addgene_125594) were derived from LRG2.1 by inserting P2A-Neo or P2A-Puro sequence after GFP sequence, respectively. For *in vitro* knockout experiments, sgRNAs targeting human or mouse gene were cloned into BsmBI-digested LRG2.1 or LRG, respectively. For *in vivo* knockout experiments, sgRNAs were cloned into BsmBI-digested LRG2.1_Neo or LRG2.1_Puro. For knockdown, shRNAs were cloned into the retroviral expression vector LEPG (Addgene_111160) according to the procedure described previously^31^. Sequences of sgRNA and shRNA are provided in Table S1. The mouse Sik3 cDNA purchased from GE Dharmacon (Clone ID: 6515742) was cloned into a LentiV_Neo vector (Addgene_108101) using In-Fusion cloning system (Clontech). The gatekeeper mutation (T142Q) was introduced by site-directed mutagenesis.

### Cell lines and virus transduction

Human and murine (RN2) AML cells^32^ were cultured in RPMI supplemented with 10% fetal bovine serum (FBS), except for OCI-AML3 and KASUMI-1 which were cultured in alpha-MEM with 20% FBS or RPMI with 20% FBS, respectively. HEK293T or NIH-3T3 cells were cultured in DMEM with 10% FBS or 10% bovine calf serum, respectively. Plat-E cells were cultured in DMEM with 10% FBS, 1 µg/ml puromicyn, and 10 µg/ml blasticidin. Penicillin/streptomycin was added to all media. All cell lines were cultured at 37°C with 5% CO2 and were periodically tested for mycoplasma and confirmed to be negative. Cas9-expressing RN2 (RN2c), NIH-3T3, and MOLM-13 cells were previously established^22,33^.

Lentivirus was produced in HEK293T cells using helper plasmids (VSVG and psPAX2 (Addgene_12260)) with Polyethylenimine (PEI 25000). Retrovirus was produced in Plat-E cells following standard procedures^34^. Target cells were spin-infected with the virus and 4 µg/mL polybrene. Media was changed at 24 hrs post-infection, and antibiotics (1-2 µg/ml puromycin, 1 mg/ml G418, or 10 µg/ml blasticidin) were added at 48 hrs post-infection if selection was needed.

### Competition-based proliferation assay

Cas9-expressing RN2 or NIH-3T3 cells were transduced with sgRNA linked with GFP in LRG vector. Percentage of GFP-positive cells was measured every 2 days using Guava EasyCyte Flow Cytometers (Millipore).

### Western blotting and fractionation

For preparation of whole cell extracts, cell pellets were suspended in Laemli sample buffer (Bio-Rad) containing 2-mercaptoethanol and boiled. The lysates were resolved in SDS-PAGE, followed by transfer to nitrocellulose membrane and immunoblotting. Fractionation was performed using a NE-PER Nuclear and Cytoplasmic Extraction Kit (Thermo Fisher Scientific) following the manufacture’s protocol. The antibodies used include HSC70, Santa Cruz Biotechnology #Sc-7298; MEF2C, Cell Signaling Technology #5030S; SIK3 (for mouse), MilliporeSigma #HPA045245; SIK3 (for human), MilliporeSigma #HPA048161; phospho-HDAC4/5/7, Cell Signaling Technology #3443; HDAC4, Cell Signaling Technology #15164 or #7628; Lamin B2, Cell Signaling Technology #12255; Actin, NeoMarkers #ACTN05; Tubulin, MilliporeSigma #T0198.

### Colony formation assay

Bone marrow cells were obtained by flushing cells from the femurs of 7- to 9-week-old female C57BL/6 mice (Taconic Biosciences) and lysed with ACK buffer (150 mM NH4Cl, 10 mM KHCO3, 0.1 mM EDTA). Normal myeloid progenitor cells were cultured in RPMI with 10% FBS, 2 ng/ml rmIL-3, 2 ng/ml rmIL-6, 10 ng/ml rmSCF and 50 µM β-mercaptoethanol. RN2 and normal myeloid progenitor cells were retrovirally transduced with shRNA in LEPG vector. On day 2 post-infection, 2,500 shRNA/GFP-positive RN2 cells or 10,000 shRNA/GFP-positive normal myeloid progenitor cells were plated into methylcellulose-based media without or with cytokines, respectively (MethoCult M3234 or MethoCult GF M3434 (Stem Cell technologies)). Cells were cultured in the media containing 2 µg/ml puromycin to select shRNA-positive cells. Blast-like (poorly differentiated) colonies were counted on day 7 post-infection (day 5 post-plating).

### SIK inhibitor treatment

Development of YKL-05-099 was previously reported^29^. For EC_50_ calculations, 1,000 cells were plated in opaque-walled 96-well plates, and mixed with serially-diluted concentrations of YKL-05-099 or 0.1% DMSO as a control. The number of viable cells was measured after 72 hrs incubation using a CellTiter Glo Luminescent Cell Viability Assay kit (Promega) with a SpectraMax plate reader (Molecular Devices) following the manufacture’s protocol. Data were analyzed using GraphPad Prism software. For Western blotting, cells were harvested after YKL-05-099 treatment at the indicated concentration and timing.

### Cell cycle and apoptosis analysis

Cell cycle analysis was performed using Vybrant DyeCycle Violet Stain (Thermo Fisher Scientific). Cells were incubated with the DNA dye at 37 °C for 2 hrs, and analyzed using a BD LSRFortessa flow cytometer (BD Biosciences) with FloJo software (TreeStar). Apoptosis was analyzed using flow cytometry with an APC Annexin V Apoptosis Detection Kit (BioLegend) following the manufacture’s protocol.

### Leukemia transplantation and *in vivo* experiments

Animal procedures and studies were conducted in accordance with the Institutional Animal Care and Use Committee (IACUC) at Cold Spring Harbor Laboratory and Dana-Farber Cancer Institute. For knockout experiments in murine AML cells *in vivo*, Cas9-expressing RN2 cells were infected with sgRNA in LRG2.1_Neo vector. sgRNA-positive cells were selected for 2 days by addition of G418, and 400,000 cells were transplanted into sub-lethally irradiated (5.5 Gy) 8-week-old female C57BL/6 mice by tail vein injection on day 3 post-infection. For knockout experiments in human AML cells *in vivo*, MV4-11 cells were transduced with luciferase and Cas9 (Blast) expression vectors, and then infected with sgRNA in LRG2.1_Puro vector. sgRNA-positive cells were selected for 3 days by the addition of puromycin, and 500,000 cells were transplanted into sub-lethally irradiated (2.5 Gy) 8-week-old female NSG mice (Jackson Laboratory) by tail vein injection on day 5 post-infection. AML development was monitored by biofluorescence imaging using an IVIS Spectrum system (Caliper Life Sciences). Images were taken 10 min after intraperitoneal injection of D-Luciferin (50 mg/kg).

For SIK inhibitor treatment in murine AML model, 500,000 RN2 cells were transplanted into sub-lethally irradiated (5.5 Gy) 8-week-old female C57BL/6 mice by tail vein injection. 6 mg/kg of YKL-05-099 was administrated via intraperitoneal injection once daily from day 1 post-transplantation for 3 weeks, and AML development was monitored by imaging as described above.

In patient-derived xenograft (PDX) model, 400,000 or 1,000,000 AML PDX cells were transplanted into 7-week-old female NSGS mice (Jackson Laboratory) for monitoring leukemia expansion or survival curve, respectively. YKL-05-099 was administrated by intraperitoneal injection at the indicated dosage from day 7 post-transplantation once daily. Leukemia burden in bone marrow was evaluated by human CD45 flow cytometry analysis 24 hours after the last dose of a 2-week treatment of YKL-05-099. Cells from mouse tissues were incubated with nonspecific binding blocker (anti-mouse/human CD16/CD32 Fcγ receptor; BD Biosciences) before staining for V450-human CD45 (BD Biosciences # 560368), and then subjected to flow cytometry analysis.

### ChIP-seq and data analysis

ChIP-seq analysis was performed essentially as previously described^22^. Briefly, MOLM-13 cells were treated with 250 nM of YKL-05-099 or 0.1% DMSO for 2 hrs, and crosslinked with 1% formaldehyde for 10 min at room temperature. Nuclear lysates were prepared by sonication (Bioruptor (Diagenode), low amplitude, On 30 s, Off 30 s, 10 cycles) and centrifugation. Supernatants were incubated with H3K27ac antibody (Abcam #ab4729) and Protein A magnetic beads (Dynabeads, Thermo) overnight for immunoprecipitation. The beads were extensively washed, and DNA was purified using a QIAquick PCR purification kit (QIAGEN) after RNase and proteinase K treatment. The ChIP-seq library was prepared using a TruSeq ChIP Sample prep kit (Illumina) following the manufacture’s protocol and analyzed by single-end sequencing using NextSeq (Illumina). H3K27ac ChIP-seq in SIK3 knockout cells and MEF2C ChIP-seq data were obtained from the previous work (GEO: GSE109492)^22^.

Bowtie2 and MACS2 were used for mapping of sequencing reads onto the reference human genome hg19 and peakcalling^35,36^. In MACS2, FDR cut off 5% with broad peak and narrow peak option was set for H3K27ac and MEF2C, respectively. BEDtools were used to merge the H3K27ac peaks in SIK3 knockout and SIK inhibitor (YKL-05-099) treatment experiments with corresponding control ChIP-seq^37^. The Bamliquidator package (https://github.com/BradnerLab/pipeline) was used for calculation of normalized tag counts. H3K27ac loci were considered as decreased when log_2_ fold-change of peak intensity was less than −1 upon knockout or treatment. TRAP web tool (http://trap.molgen.mpg.de/cgi-bin/home.cgi) was used for analysis of transcription factor binding motifs within 500 bp around the center of the decreased H3K27ac regions^38^. The UCSC genome browser was used to generate ChIP-seq tracks^39^. Metagene plot and heatmap density plots were generated using ±5 kb around each center of the decreased H3K27ac peaks or the same number of random H3K27ac peaks with 50 bp binning size.

### RNA-seq and data analysis

RNA-seq analysis was performed as previously described with some modifications^22^. Briefly, cells were harvested after the indicated concentration of YKL-05-099 for 2 hrs, or on 2 days post-infection of sgRNA for gene knockout in RN2 cells. The RNA-seq library was prepared using 2 µg of total RNA with a TruSeq sample prep kit v2 (Illumina) according to the manufacture’s protocol and analyzed by single-end sequencing using NextSeq (Illumina).

HISAT2 was used to map sequencing reads to the reference human (hg19) or mouse (mm10) genome^40^. FeatureCounts was used to count the reads assigned to protein coding genes^41^. DESeq2 was used for comparison of gene expressions^42^. Custom gene signatures for Gene Set Enrichment Analysis (GSEA) were prepared from top 200 downregulated genes upon gene knockout or YKL-05-099 treatment. Gene signatures for SIK3 or MEF2C knockout in MOLM-13 were obtained from the previous work (GEO: GSE109491)^22^ GSEA was performed according to the instructions using these custom signatures^43^.

ChIP-seq and RNA-seq data prepared in this paper are available in NCBI’s Gene Expression Omnibus database^44^ through GEO Series accession number GSE129863.

## Results

### SIK3 and MEF2C are selectively essential for the growth of acute myeloid leukemia and multiple myeloma cells

To complement our prior mechanistic characterization of the SIK3-MEF2C co-dependency in AML^22^ (Fig. S1A), we explored here the therapeutic significance of SIK3 with an expanded genetic and pharmacological evaluation of this target. Since our prior CRISPR screens were limited to 26 cancer cell lines, we began by examining the essentiality of SIK3 in a larger collection of cancer contexts using recently published genome-wide CRISPR screening datasets^45^. Across more than 500 lines representing 28 different cancer cell lineages, we found that SIK3 was exclusively essential in AML and multiple myeloma contexts (Fig. 1A and S1B). Importantly, this pattern resembled that of MEF2C, which was also selectively essential in these two cancer subtypes (Fig. 1A and S1C). A correlation between SIK3 and MEF2C dependence in AML was observed in these datasets (Fig. 1B) and also in an independent set of CRISPR screens performed by Wang & Sabatini et al^46^ (Fig. 1C). The pattern of SIK3 essentiality is distinct from other kinases, such as CDK1, which lacks specificity for any particular tumor subtype (Fig. S1B). Notably, multiple myeloma and AML cell lines express the highest levels of MEF2C and SIK3 when compared to other cancer cell lines^47^ (Fig. S1D). Together, these findings suggest that SIK3 and MEF2C are not general requirements for cell proliferation, but their essentiality is highly specific to the hematological malignancies AML and multiple myeloma.

**Figure 1.**
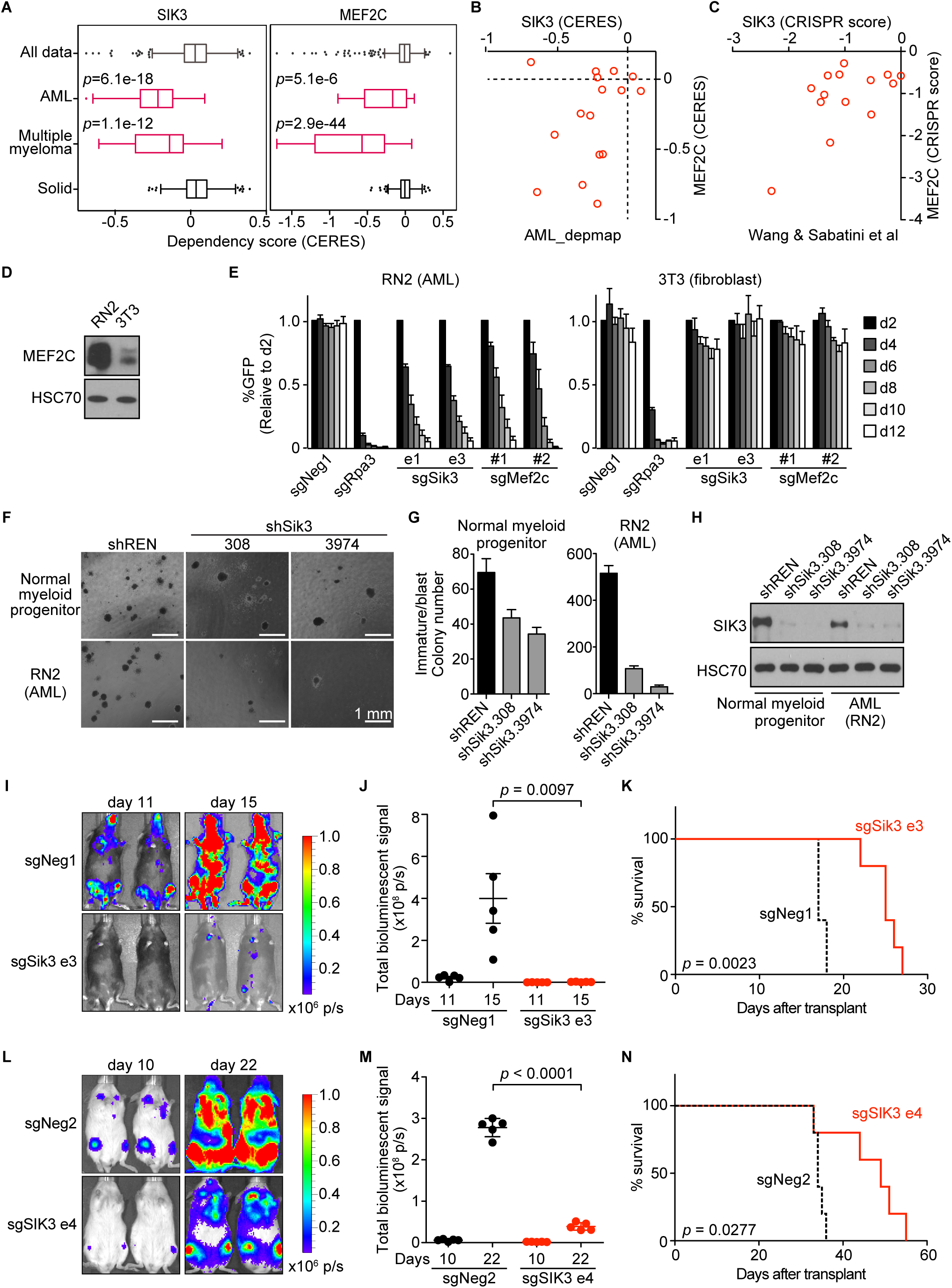
SIK3 and MEF2C are selectively essential for the growth of acute myeloid leukemia and multiple myeloma cells. (A) SIK3 and MEF2C essentiality scores extracted from the DepMap database (19Q1) of cancer cell lines^45^. Shown is a boxplot distribution of CERES scores (a normalized metric of gene essentiality) of SIK3 and MEF2C across all 558 cell lines, 16 AML lines, 18 multiple myeloma lines, or 498 solid tumor cell lines. (B-C) Scatterplots of SIK3 and MEF2C essentiality scores in human AML cell lines in the DepMap (CERES) or from Wang & Sabatini. et al.^46^ (CRISPR scores). (D) Western blot analysis of MEF2C in RN2 and 3T3 whole cell lysates. (E) Competition-based proliferation assays in which cells were infected with the indicated sgRNAs linked to GFP. Bar graphs represent the mean ± SEM (n=3). (F) Bright-field images of methylcellulose-based colony formation assays of normal myeloid progenitors or RN2 cells on day 7 post retroviral transduction with control or Sik3 shRNAs. (G) Quantification of immature/blast colonies shown in (F). Mean ± SEM (n = 4). (H) Western blot analysis of SIK3 performed on day 6 post-infection of the indicated shRNAs. (I) Bioluminescence imaging of wild-type C57BL/6 mice transplanted with Cas9-expressing RN2 cells transduced with the indicated sgRNA. Representative images are shown on the indicated day post-transplantation. (J) Quantification of bioluminescence from (I). Values represent photons per second (p/s) of bioluminescent signal detection. The *p*-value was calculated using unpaired Student’s t test (n = 5). (K) Survival curves of mice used in (I). The *p*-value was calculated using log rank (Mantel-Cox) test (n = 5). (L) Bioluminescence imaging of NSG (NOD-scid/IL2Rgamma^null^) mice transplanted with Cas9-expressing MV4-11 cells transduced with the indicated sgRNA. Representative images are shown on the indicated day post-transplantation. (M) Quantification of bioluminescence from (L). Values represent photons per second (p/s) of bioluminescent signal detection. The *p*-value was calculated using unpaired Student’s t test (n = 5). (N) Survival curves of mice used in (L). The *p*-value was calculated using log rank (Mantel-Cox) test (n = 5). sgNeg1, sgNeg2, shREN: negative controls. sgRpa3: positive control

To further validate the specificity of SIK3 and MEF2C as dependencies in AML, we inactivated each gene in immortalized 3T3 murine fibroblasts and in the murine AML cell line RN2, which was generated by expressing the MLL-AF9 and Nras^G12D^ oncogenes in hematopoietic stem and progenitor cells^32^. As expected, RN2 cells express higher levels of MEF2C than 3T3 cells (Fig. 1D). Using CRISPR-Cas9 genome editing and competition-based assays of cellular fitness, we found that inactivating *Sik3* or *Mef2c* elicited a severe growth arrest in RN2 cells while the same manipulation had no detectable impact on the growth of 3T3 cells (Fig. 1E and S2A). We further compared the specificity of Sik3 dependence by using short hairpin RNAs to acutely deplete Sik3 in RN2 and in bone marrow-derived myeloid progenitor cells. After plating in methylcellulose, we observed a stronger reduction of myeloid colony formation upon knockdown of Sik3 in RN2 than in normal myeloid cells (Fig. 1F-H). These findings are in agreement with the prior characterization of *Sik3* and *Mef2c* knockout mice, which lack a defect in normal myelopoiesis^12,48^. These findings also corroborate our prior analysis of human AML cell lines^22^, and suggest an enhanced dependency on SIK3 and MEF2C in *MLL*-rearranged leukemia relative to normal myeloid cells.

We next investigated whether SIK3-deficient AML cells exhibit a growth defect under *in vivo* conditions. To this end, control and *Sik3* knockout RN2 cells were injected via tail vein into sub-lethally irradiated C57BL/6 mice. AML progression was monitored using bioluminescent imaging of luciferase, which is constitutively expressed from a retroviral promoter in RN2 cells^32^ (Fig. S2B-C). Notably, inactivation of *Sik3* led to a marked attenuation of AML progression and extended animal survival (Fig. 1I-K). Similar results were obtained upon transplantation of *SIK3* knockout MV4-11 cells (a human MLL-AF4 AML line) into immune-deficient NSG mice (Fig. 1L-N and S2D-E). These findings demonstrate that SIK3 is essential for the expansion of MLL fusion AML cells under both *in vitro* and *in vivo* conditions.

### YKL-05-099 inhibits the growth of *MLL*-rearranged leukemia cells by modulating SIK3-mediated regulation of HDAC4

The specificity and *in vivo* relevance of SIK3 essentiality in AML led us to investigate pharmacological approaches to inhibit SIK3 as a therapeutic strategy. The tool compound HG-9-91-01 is a widely used pan-SIK inhibitor for tissue culture studies^49^, which we previously used to validate SIK3 as a dependency in AML^22^. However, HG-9-91-01 is rapidly degraded in liver microsomes and is not suitable for *in vivo* studies^29^. To overcome this issue, we recently developed YKL-05-099, which has improved pharmacokinetic properties relative to HG-9-91-01 (Fig. 2A)^29,50^. Using comparative kinome profiling, we have found that YKL-05-099 also exhibits improved selectivity for SIKs over other kinases and a modest decrease in potency relative to HG-9-91-01^29^.

**Figure 2.**
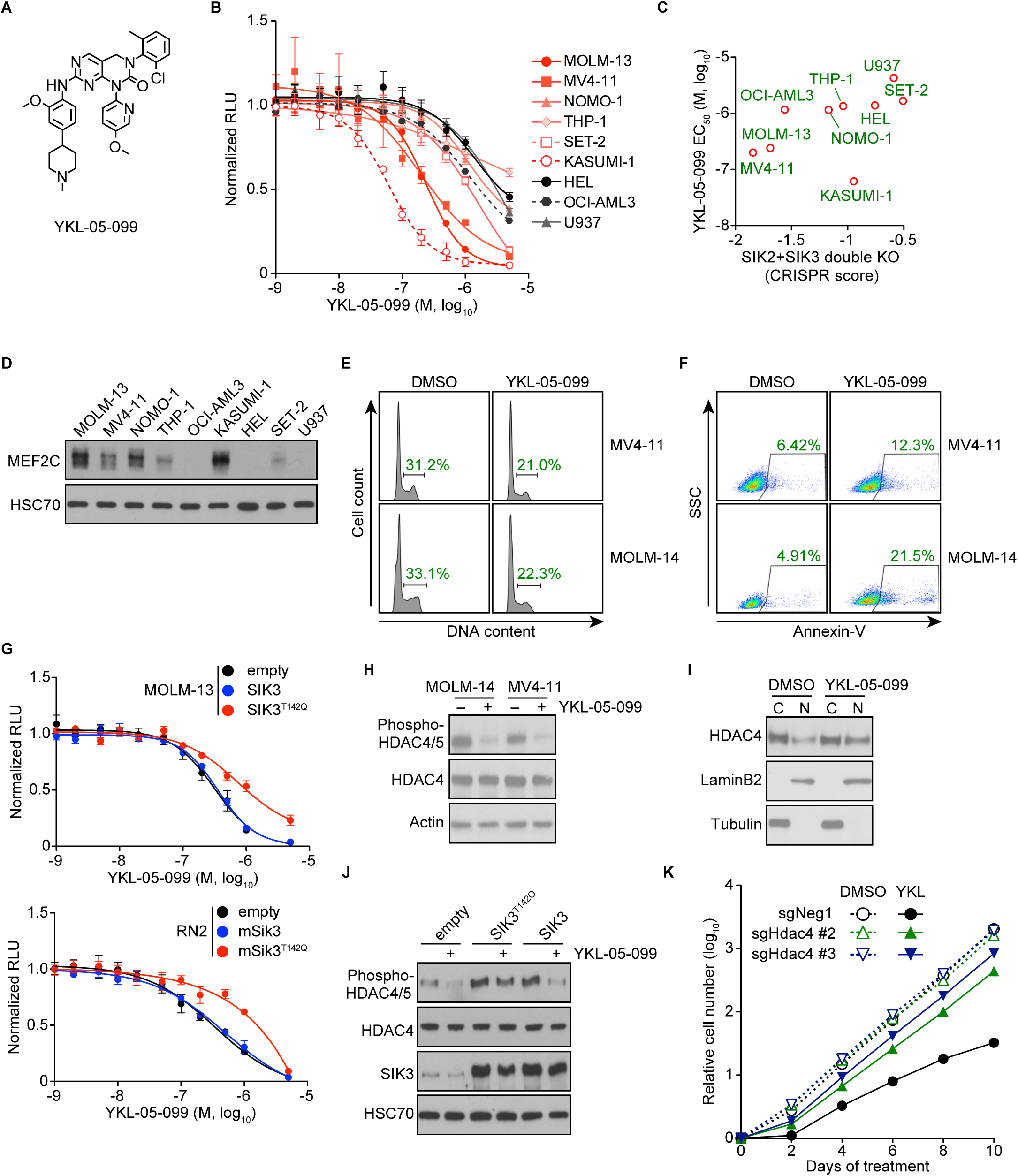
YKL-05-099 inhibits the growth of *MLL*-rearranged leukemia cells by modulating SIK3-mediated regulation of HDAC4. (A) Chemical structure of YKL-05-099. (B) Relative growth of the indicated cells using CellTiter-Glo assays. Normalized relative luminescence unit (RLU) is shown after 3 days in culture with DMSO (0.1%) or YKL-5-099 at the indicated concentrations. The mean ± SEM (n=3) and four-parameter dose-response curves are plotted. (C) Scatterplot of SIK2/SIK3 essentiality CRISPR score from our prior study^22^ and YKL-05-099 EC_50_ in the indicated AML cell lines. The CRISPR scores were calculated in cells co-transduced with SIK2 and SIK3 sgRNAs and cell fitness tracked in competition-based assays. (D) Western blot analysis of MEF2C in the indicated AML cell lines. (E) Flow cytometry analyses of DNA content to infer cell cycle status after 24 hour treatment with 1 µM of YKL-05-099 or vehicle (DMSO) control. (F) Flow cytometry analyses of side scatter and Annexin-V staining (a pre-apoptotic cell marker) after 24 hour treatment with 1 µM of YKL-05-099 or vehicle (DMSO) control. (G) Relative growth of RN2 and MOLM-13 cells transduced with empty vector, *SIK3*, or *SIK3*^T142Q^ cDNA, upon YKL-05-099 treatment. Normalized relative luminescence unit (RLU) is shown after 3 days culture with DMSO (0.1%) or YKL-05-099 at the indicated concentrations. The mean ± SEM (n=3) and four-parameter dose-response curves are plotted. (H) Western blot analysis in MOLM-14 and MV4-11 cells treated with 1 µM of YKL-05-099 or vehicle for 6 hrs. (I) Western blot analysis of fractionated proteins in MV4-11 cells after treatment with 1 µM of YKL-05-099 for 24 hrs. C, cytosol; N, nucleus. (J) Western blot analysis in RN2 cells transduced with empty vector, *Sik3*, or *Sik3*^T142Q^ cDNA, following treatment with DMSO (0.1%) or 350 nM of YKL-05-099 for 2 hrs. (K) Accumulated cell number of Cas9-expressing RN2 cells transduced with the indicated sgRNAs upon treatment with DMSO (0.1%) or 350 nM of YKL-05-099. Average of three independent experiments is shown. sgNeg1: negative controls.

Using a 12-point compound titration and CellTiter-Glo proliferation assays, we found that the growth of AML cell lines was inhibited by YKL-05-099 with an (e.g., an EC50 of 240 nM in MOLM-13 cells) in a manner that correlated with the overall sensitivity to *MEF2C* and *SIK2/SIK3* knockout^22^ (Fig. 2B-C and S3A). In addition, Western blot analysis showed that AML cells with high MEF2C protein expression tended to be more sensitive to YKL-050-99 (Fig. 2D). Flow cytometry analysis revealed that YKL-05-099 treatment led to a G1/G0 cell cycle arrest and the induction of apoptosis in *MLL*-rearranged human AML lines (Fig. 2E-F). Since YKL-05-099 inhibits several cancer-relevant kinases in addition to SIKs (e.g., SRC and ABL) (Fig. S3B)^29^, we also pursued genetic confirmation that YKL-05-099-induced growth arrest occurred via an on-target inhibition of SIK3. For this purpose, we made use of a gatekeeper mutant SIK3^T142Q^, which has been shown previously to render this kinase resistant to chemical inhibition^22^. Importantly, lentiviral expression of SIK3^T142Q^, but not wild-type SIK3, alleviated the sensitivity to YKL-05-099-mediated growth arrest in both RN2 and MOLM-13 cell contexts (Fig. 2G). This rescue experiment indicates that on-target SIK3 inhibition contributes to the growth arrest caused by YKL-05-099 in AML.

We previously showed that the critical substrate of SIK3 in supporting AML growth is HDAC4^22^, which is sequestered in the cytoplasm by SIK3-mediated phosphorylation^27^ (Fig. S1A). Based on these prior observations, we examined whether the YKL-05-099 compound would also modulate HDAC4 function in AML. Western blotting analysis revealed that YKL-05-099 treatment led to rapid dephosphorylation and nuclear accumulation of HDAC4 in AML cells (Fig. 2H-I and Fig. S3C). Importantly, this effect occurred via inhibition of SIK3, since lentiviral expression of the SIK3^T142Q^ cDNA rescued this effect (Fig. 2J). We next used CRISPR-Cas9 to inactivate HDAC4 in RN2 and MOLM-13 cells (Fig. S3D). While the knockout of HDAC4 did not influence the growth rate of these lines, the lack of HDAC4 rendered both AML contexts resistant to YKL-05-099-mediated growth arrest (Fig. 2K and Fig. S3E). These experiments lend further support that YKL-05-099 causes growth arrest in *MLL*-rearranged leukemia cells by releasing HDAC4 from SIK3-mediated sequestration.

### YKL-05-099 interferes with MEF2C-dependent transcriptional activation

Our prior epigenomic analysis of MOLM-13 cells revealed that *SIK3* knockout led to the selective loss of histone acetylation at enhancer elements occupied by MEF2C and a reduced mRNA level of MEF2C target genes^22^. To complete our drug mechanism-of-action study, we investigated whether YKL-05-099 treatment would also lead to similar changes in acetylation and transcription. To this end, we performed chromatin immunoprecipitation of histone H3 lysine 27 acetylation (H3K27ac, a mark of active promoters and enhancers) and deep sequencing (ChIP-seq) following a 2-hr treatment of MOLM-13 cells with 250 nM YKL-05-099 or with vehicle control. By comparing these two acetylation landscapes, we found that H3K27ac was selectively reduced at select genomic sites (Fig. 3A-B). Importantly, YKL-05-099-induced losses of H3K27ac correlated with acetylation changes observed following *SIK3* knockout and were prevented by SIK3^T142Q^ expression, suggesting they occurred via on-target SIK3 inhibition (Fig. 3B-C). In addition, a sequence analysis of these genomic intervals revealed an enrichment of MEF2C motifs at these locations (Fig. 3D). Moreover, our prior ChIP-seq datasets confirmed that MEF2C occupancy was found at 95% of the *cis*-elements with reduced H3K27ac following YKL-05-099 exposure^22^ (Fig.3A, E-F).

**Figure 3.**
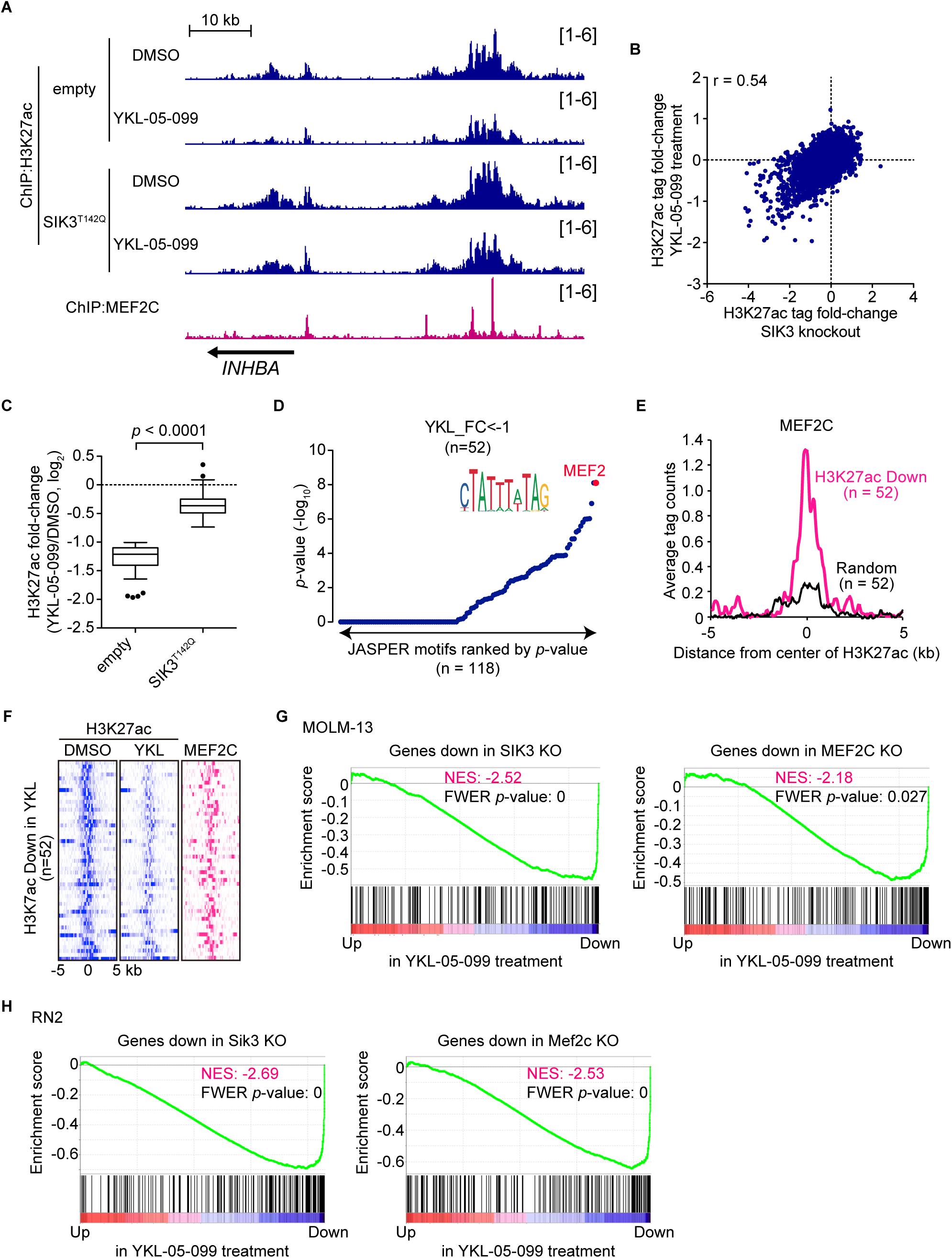
YKL-05-099 interferes with MEF2C-dependent transcriptional activation. (A) ChIP-seq profiles of H3K27ac and MEF2C at the indicated genomic loci, chosen because H3K27ac is decreased upon YKL-05-099 treatment in MOLM-13 cells. For H3K27ac ChIP-seq, cells were harvested after 2-hrs treatment with DMSO (0.1%) or 250 nM YKL-05-099. (B) Scatterplot of H3K27ac fold-change upon *SIK3* knockout or YKL-05-099 treatment at 16,437 genomic sites of H3K27ac enrichment. (C) Box plots of fold-change of down-regulated H3K27ac signals in MOLM-13 cells treated with DMSO or 250 nM of YKL-05-099 for 2 hrs. MOLM-13 cells express either an empty vector or *SIK3*^T142Q^ cDNA. (D) TRAP motif enrichment analysis of DNA sequences with decreased H3K27ac upon YKL-05-099 treatment. (E) A meta-profile of MEF2C occupancy at the genomic regions exhibiting H3K27ac log2 fold-change of <-1 after YKL-05-099 treatment, versus a randomly chosen set of H3K27ac-enriched sites. (F) ChIP-seq density plot at regions with decreased H3K27ac upon YKL-05-099 treatment. Enhancers are ranked by fold-change of H3K27ac upon YKL-05-099 treatment. (G) Genes Set Enrichment Analysis (GSEA) that evaluates how treating MOLM-13 cells with YKL-05-099 (250 nM, 2 hrs) influences previously defined gene signatures are suppressed following *SIK3* or *MEF2C* knockout in this cell type^22^. Normalized enrichment score (NES) and family-wise error rate (FWER) *p*-value are shown. (H) Genes Set Enrichment Analysis (GSEA) that evaluates how treating RN2 cells with YKL-05-099 (250 nM, 2 hrs) influences gene signatures that are suppressed following *Sik3* or *Mef2c* knockout in this cell type. Normalized enrichment score (NES) and family-wise error rate (FWER) *p*-value are shown.

We also performed RNA-seq analysis of MOLM-13 cells following a 2 hour exposure to YKL-05-099 and compared the results to our prior RNA-seq analysis of *SIK3* or *MEF2C* knockout in this same cellular context^22^. Using Gene Set Enrichment Analysis^43^, we found YKL-05-099 treatment rapidly suppressed a similar transcriptional program as observed following genetic targeting of SIK3 or MEF2C (Fig. 3G). To further corroborate this finding, we performed a similar RNA-seq evaluation in the RN2 cell line after *Sik3* knockout, *Mef2c* knockout, and YKL-05-099 exposure. These experiments revealed a significant overlap of transcriptional changes following all three perturbations (Fig. 3H). However, we note across all of our experiments that a genetic knockout of *SIK3* tended to have a stronger effect on H3K27ac enrichment than YKL-05-099 treatment (Fig. 3B). Nevertheless, these findings lend strong support for on-target SIK3 inhibition via YKL-05-099, thus validating the use of this compound as a pharmacological strategy to suppress the transcriptional output of MEF2C in AML.

### YKL-05-099 treatment extends survival in two mouse models of MLL-AF9 AML

Having demonstrated the activity of YKL-05-099 in interfering with the SIK3-HDAC4-MEF2C signaling axis, we next evaluated the efficacy of this agent in animal models of *MLL*-rearranged AML. To this end, we transplanted RN2 cells into sub-lethally irradiated C57BL/6 mice, followed by administration of YKL-05-099 by intraperitoneal injection for three weeks, starting on day 1 post-transplantation (Fig. 4A-B). Bioluminescence imaging of luciferase revealed a significant delay in AML progression, which was associated with a survival benefit in the YKL-05-099 treated cohort (*p* = 0.0027) (Fig. 4B-D). We further investigated the efficacy of YKL-05-099 in a patient-derived xenograft (PDX) model of MLL-AF9 AML propagated in NSGS mice. Flow cytometry analysis of bone marrow cells after 2 weeks of YKL-05-099 treatment in the PDX model revealed a dose-dependent decrease in human AML cells (Fig. 4E). Importantly, all three doses of YKL-05-099 did not significantly reduce the weight of mice during this 2-week period of treatment (Fig. 4F). In an independent cohort of AML PDX-bearing mice, a 3-week treatment with YKL-05-099 was associated with a significant extension of animal survival (*p* < 0.001) (Fig. 4G). Taken together, these experiments suggest that YKL-05-099 treatment can suppress AML progression and extend animal survival at well-tolerated doses.

**Figure 4.**
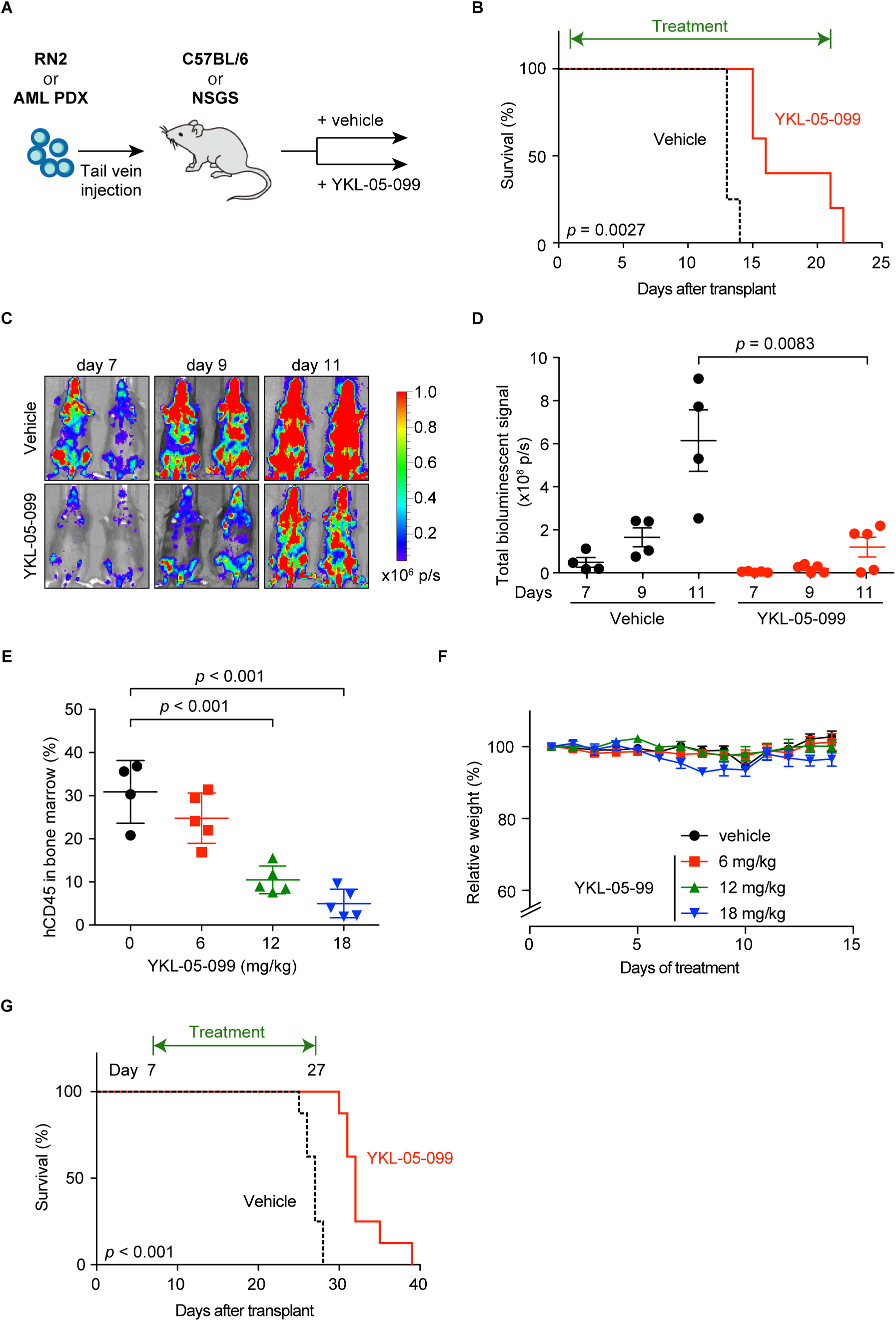
YKL-05-099 treatment extends survival in two mouse models of MLL-AF9 AML. (A) Diagram of YKL-05-099 treatment experiments in AML mouse models. (B) Survival curves of C57BL/6 mice transplanted with RN2 cells, followed by intraperitoneal injection of 6 mg/kg YKL-05-099 treatment once daily from day 1 post-transplantation. The *p*-value was calculated using log rank (Mantel-Cox) test (n = 4 or 5). (C) Representative bioluminescence imaging of mice used in (B) on the indicated days post-transplantation. (D) Quantification of signal from (C). Values represent photons per second (p/s) of bioluminescent signal detection. The *p*-value was calculated using unpaired Student’s t test (n = 4 or 5). (E) Leukemia burden in bone marrow was evaluated by human CD45 flow cytometry analysis after 2-weeks treatment of YKL-05-099 in NSGS mice transplanted with AML PDX model. The mean ± SD is shown (n = 4 or 5). The *p*-value was calculated using unpaired Student’s t test. (F) Mouse weight measurements performed during YKL-05-099 treatment from in (E). (G) Survival curves of NSGS mice transplanted with AML PDX cells. YKL-05-099 treatment (18 mg/kg, intraperitoneal injection, once daily) was initiated from day 7 post-transplantation for three weeks. The *p*-value was calculated using log rank (Mantel-Cox) test (n = 8).

## Discussion

The goal of our study was to evaluate the therapeutic potential of targeting the SIK3-HDAC4-MEF2C signaling axis in AML. Using a collection of genetic and pharmacological experiments, our work suggests that a therapeutic window exists when targeting SIK3 in the *MLL*-rearranged subtype of AML. In addition, our findings suggest that SIK3 will have relevance as a target in other malignancies that are addicted to MEF2C, such as multiple myeloma. The unique attribute of SIK3 as a target is the selectivity of its essentiality, which is now supported by CRISPR screening results from over 500 cancer cell lines and by our experiments in normal hematopoietic cells. Unlike other non-oncogene kinase targets (e.g. CDK1), we expect that selective inhibitors of SIK3 will have minimal on-target effects on the growth of non-malignant tissues, which is supported by the limited toxicities observed in YKL-05-099-treated mice and by the viability of *Sik3*-deficient mice^48^. It is likely that other kinases (e.g., CaMK and AMPK) compensate for the loss of SIK3 in various normal tissues to maintain HDAC4 phosphorylation^23,51^, which may account for the tolerance of mice to sustained SIK inhibition.

One caveat of our animal studies is that YKL-05-099 treatment only led to a modest attenuation of AML progression. However, we note throughout our study that genetic inactivation of SIK3 leads to stronger effects on cell growth, histone acetylation, and gene expression than our highest on-target doses of YKL-05-099. This suggests that efficacy in our animal studies is limited by the pharmacological properties of YKL-05-099, rather than a limitation of SIK3 as a genetic dependency. While our chemical optimization of YKL-05-099 has led to improved pharmacokinetic properties^29^ (e.g. free serum concentrations of YKL-05-099 exceed the IC_50_ for SIK3 inhibition for > 16 hr *in vivo* after intraperitoneal injection), the overall potency of this compound and its selectivity for SIKs will require further optimization to reach the profile of other successful kinase inhibitors used in the clinic. The docking study of YKL-05-099 into a SIK3 homology model suggests several possible focus points for further medicinal chemistry efforts. In parallel, an additional lead has been identified from scaffold-morphing and chimeric analogs will be explored by combining structural feature of these two leads. Additionally, small-molecule degraders, also referred to as proteolysis targeting chimeras or degronimids, could potentially improve the kinome selectivity and enable abrogation of non-kinase dependent function of SIKs as a strategy to obtain maximal MEF2C suppression in AML inhibitors^52–55^. Our medicinal chemistry efforts are now ongoing to advance a drug-like compound for definitive pre-clinical investigation of SIK inhibition as a therapeutic approach in AML, as well as in other hematopoietic cancers with MEF2C addiction.

The key mechanistic advance in our study is in reinforcing the potential of lineage TFs as therapeutic targets in AML, which we have shown can be chemically disabled by targeting of upstream signaling pathways^56^. Genetic studies by our lab and others have identified MYB, PU.1, CEBPA, RUNX1, and MEF2C as elite non-oncogene dependencies in AML, both in terms of potency and selectivity of their essential function^22,57–61^. From this perspective, an approach that seeks to define upstream kinase signals that support each of these essential TFs may provide a source of novel AML therapies.

## Acknowledgments

This work was supported by Cold Spring Harbor Laboratory NCI Cancer Center Support grant 5P30CA045508. Additional funding was provided by the Forbeck Foundation, Pershing Square Sohn Cancer Research Alliance, National Institutes of Health grants R35 CA210030, R01 CA174793 and P01 CA013106, and a Leukemia & Lymphoma Society Scholar Award. Y.T. was supported by the Lauri Strauss Leukemia Foundation.

## Authorship

Y.T., S.L., J.W., J.P.M., and Y.X. performed experiments; Y.T. and S.L. analyzed the data; N.S.G, K.S., and C.R.V. supervised the experiments and analysis; Y.T., S.L., K.S., and C.R.V. wrote the manuscript.

---

**Figure S1.**
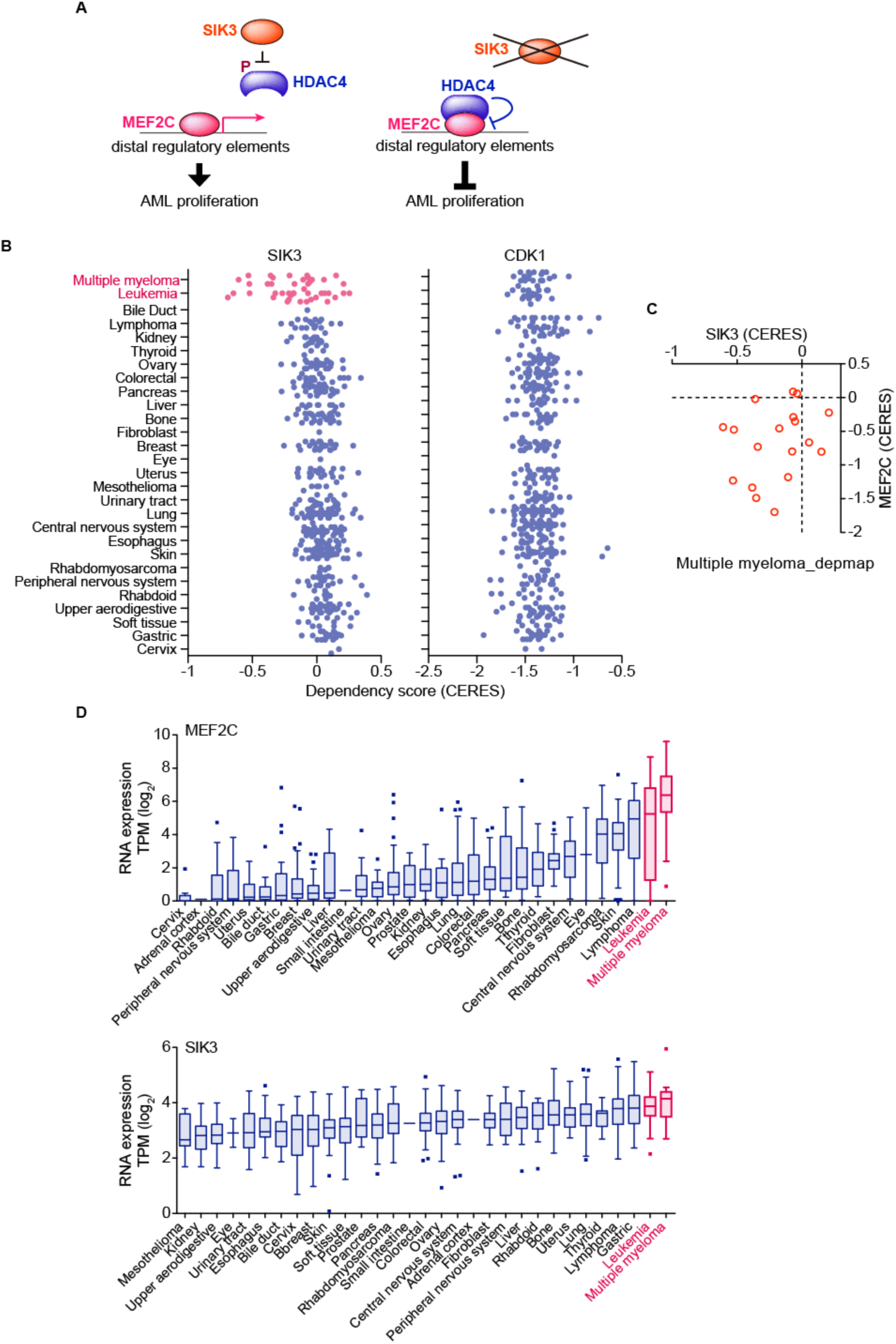
SIK3 is selectively essential for growth of acute myeloid leukemia and multiple myeloma. (A) Schematic diagram depicting the regulation of MEF2C function by SIK3 through HDAC4 phosphorylation. (B) SIK3 and CDK1 essentiality scores extracted from the DepMap database (19Q1) of cancer cell lines^46^. Shown is a boxplot distribution of CERES scores across each cancer lineage. (C) Scatterplots of SIK3 and MEF2C essentiality scores in human multiple myeloma cell lines in the DepMap (CERES) (D) SIK3 and MEF2C RNA expression level of MEF2C and SIK3 extracted from the CCLE database^48^. Shown is a boxplot distribution of TPM (Transcripts Per Kilobase Million) across each cancer lineage.

**Figure S2.**
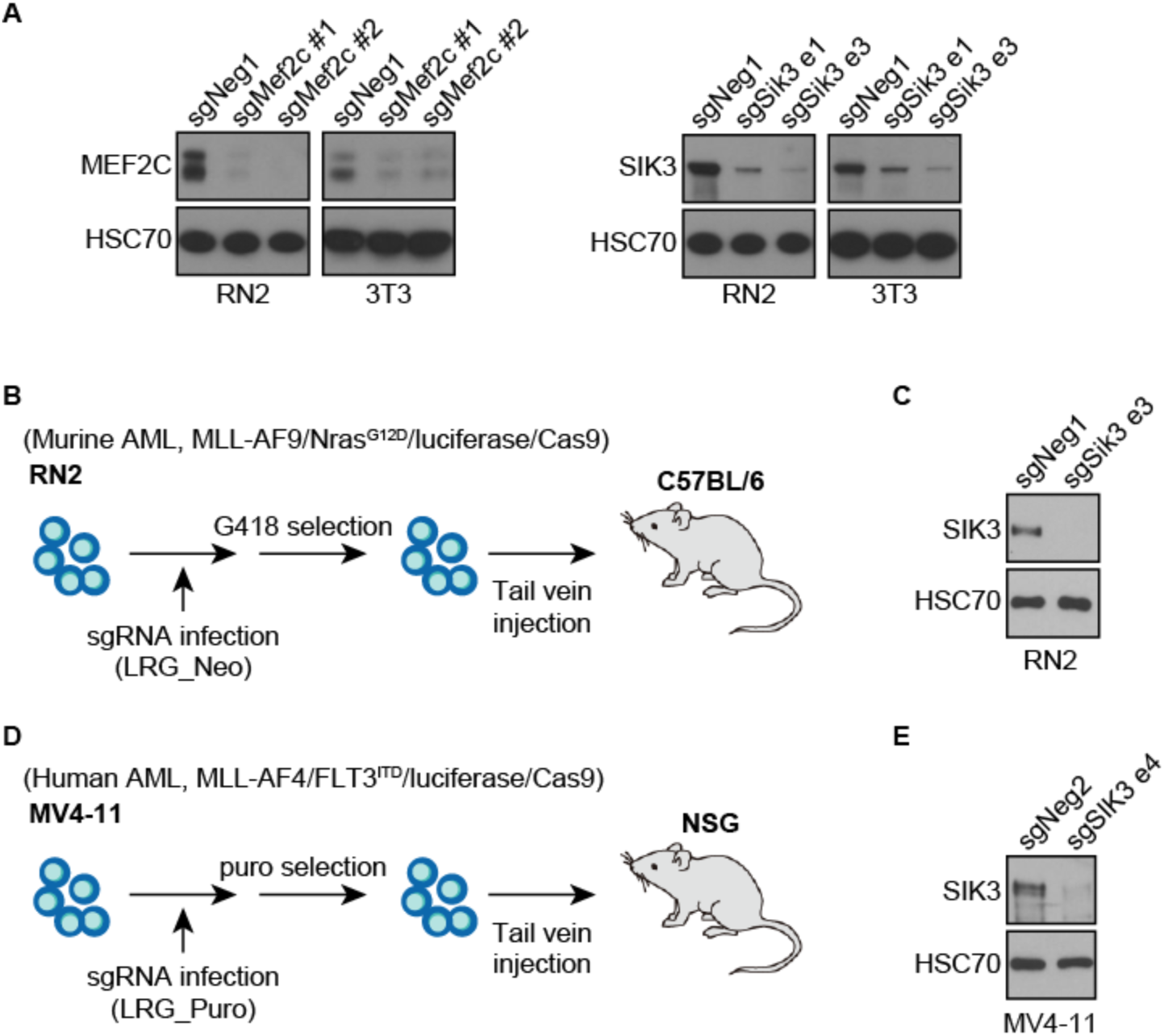
SIK3 is important for AML growth *in vitro* and *in vivo*. (A) Western blot analysis of SIK3 and MEF2C in RN2 and 3T3 cells on day 2 post-infection with the indicated sgRNAs. (B and D) Schematic diagram depicting transplantation of AML cells into mice. (C and E) Western blot analysis of SIK3 in the cells on day 3 (C) or day 5 (E) post-infection with the indicated sgRNA. sgNeg1, sgNeg2: negative controls.

**Figure S3.**
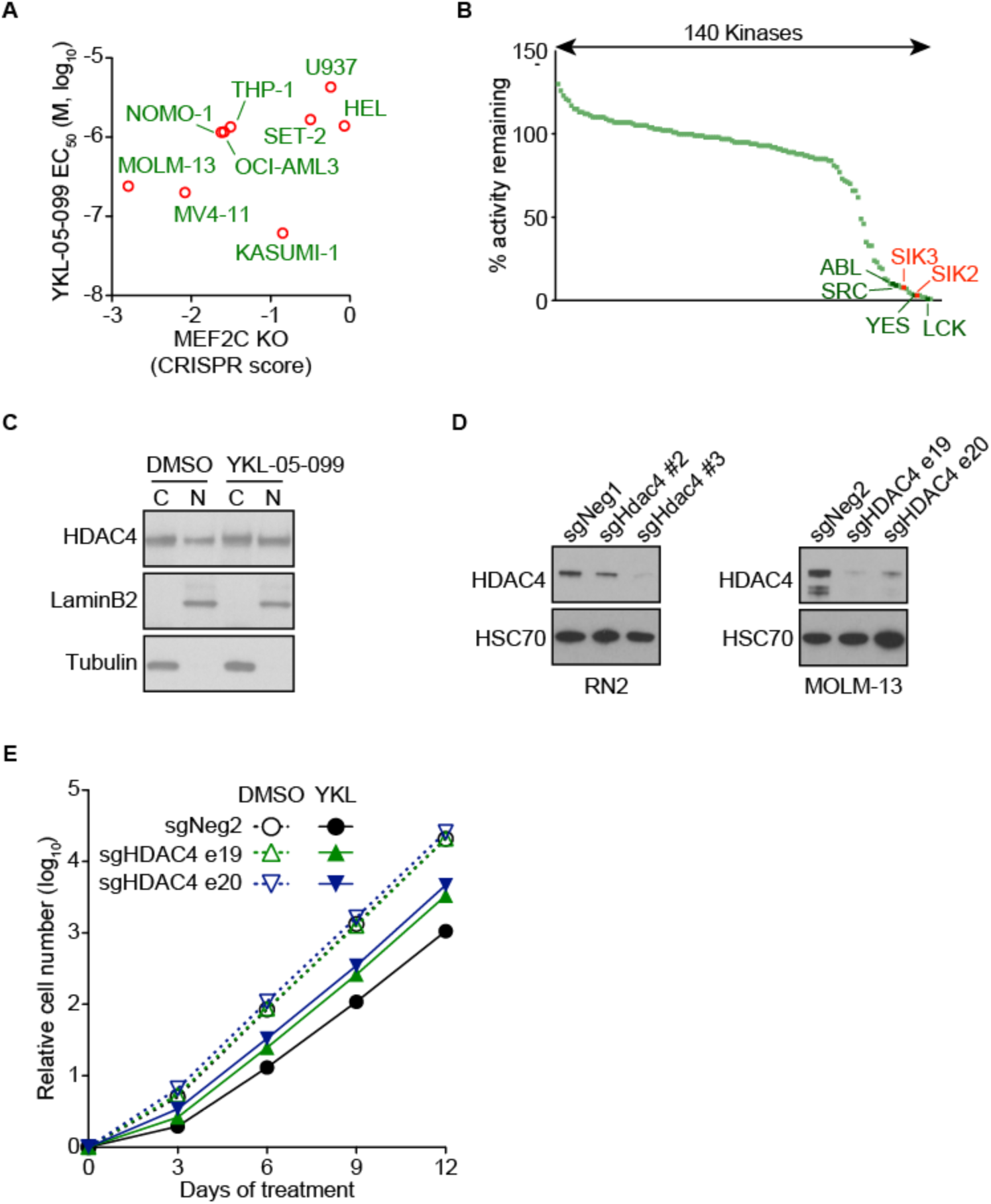
YKL-05-099 inhibits AML growth by modulating SIK3-mediated regulation of HDAC4. (A) Scatterplot of MEF2C essentiality CRISPR score from our prior study^22^ and YKL-05-099 EC_50_ in the indicated AML cell lines. The CRISPR scores were calculated in cells transduced with MEF2C sgRNAs and cell fitness tracked in competition-based assays. (B) The *in vitro* activity of 140 kinases in the presence of 1 µM of YKL-05-099 extracted from Sundberg et al.^29^ (C) Western blot analysis of fractionated proteins in MOLM-14 cells after treatment of 1 µM of YKL-05-099 for 24 hrs. C, cytosol; N, nucleus. (D) Western blot analysis of HDAC4 in Cas9-expressing RN2 and MOLM-13 cells on days 2 and 3 post-infection with the indicated sgRNAs, respectively. (E) Growth curves of Cas9-expressing MOLM-13 cells infected with the indicated sgRNA upon treatment with DMSO (0.1%) or 250 nM of YKL-05-099. Average of three independent experiments are shown. sgNeg1 and sgNeg2: negative controls.

